# Integrating cyanobacterial flavodiiron proteins within the chloroplast photosynthetic electron transport chain maintains carbohydrate turnover and enhances drought stress tolerance in barley

**DOI:** 10.1101/2020.09.29.318394

**Authors:** Fahimeh Shahinnia, Suresh Tula, Goetz Hensel, Narges Reiahisamani, Nasrin Nasr, Jochen Kumlehn, Rodrigo Gómez, Anabella F. Lodeyro, Néstor Carrillo, Mohammad R. Hajirezaei

**Affiliations:** Department of Physiology and Cell Biology, Leibniz Institute of Plant Genetics and Crop Plant Research, OT Gatersleben, Corrensstrasse 3, D-06466 Seeland, Germany; Division of Molecular Biology, Centre of the Region Hana for Biotechnological and Agriculture Research, Faculty of Science, Palacký University, Olomouc, Czech Republic; Centre of Plant Genome Engineering, Institute of Plant Biochemistry, Heinrich-Heine-University, Dusseldorf, Germany; Instituto de Biología Molecular y Celular de Rosario (IBR-UNR/CONICET), Facultad de Ciencias Bioquímicas y Farmacéuticas, Universidad Nacional de Rosario, Rosario, Argentina; Dipartimento di Biotecnologie, Università di Verona, Strada Le Grazie 15, 37134 Verona, Italy

**Author notes:** **Correspondence:** Dr. Mohammad-Reza Hajirezaei.

**Keywords:** Biomass, *Hordeum vulgare* L., Metabolites, Photosynthesis, Plastid biotechnology, Yield

## Abstract

Chloroplasts, the sites of photosynthesis in higher plants, have evolved several means to tolerate short episodes of drought stress through biosynthesis of diverse metabolites essential for plant function, but these become ineffective when the duration of the stress is prolonged. Cyanobacteria are the closest bacterial homologs of plastids with two photosystems to perform photosynthesis and to evolve oxygen as a byproduct. The presence of *Flv* genes encoding flavodiiron proteins has been shown to enhance stress tolerance in cyanobacteria. Here, the products of *Synechocystis* genes *Flv1* and *Flv3* were expressed in chloroplasts of barley in an attempt to support the growth of plants exposed to drought. The heterologous expression of both *Flv1 and Flv3* accelerated days to heading, increased biomass, promoted the number of spikes and grains per plant, and improved grain yield of barley plants exposed to drought. Improved growth correlated with enhanced availability of soluble sugars, a higher turnover of amino acids and the accumulation of lower levels of proline in the leaf. *Flv1* and *Flv3* maintained the energy status of the leaves in the stressed plants by converting sucrose to glucose and fructose, immediate precursors for energy production to support plant growth under drought. The results suggest that sugars and amino acids play a fundamental role in the maintenance of the energy status and metabolic activity to ensure growth and survival under stress conditions, that is, water limitation in this particular case. Engineering chloroplasts by introducing *Flv* genes, therefore, has the potential to improve plant productivity wherever drought stress represents a significant production constraint.

## INTRODUCTION

If current predictions indicating that the world’s population will rise to 9.4 billion by 2050 prove correct (Wang et al., 2013), a substantial increase in crop production will be required to meet the global demand for food. However, levels of crop productivity are unlikely to keep pace with this demand (Ray et al., 2012, 2013), without technological interventions directed to enhance photosynthetic efficiency and/or bolster tolerance to abiotic stress (Cardona et al., 2018; Gómez et al., 2019; Batista-Silva et al., 2020). Drought poses a major constraint over crop productivity (Wang et al., 2011), both directly and through its aggravation of the impact of other stress factors.

To date, biochemical studies have been extensively used to elucidate the metabolic responses to abiotic stresses (especially drought) in different plant species and to improve stress tolerance (Fàbregas and Fernie, 2019, and references therein). In general, plants respond to water restriction by closing their stomata, which in turn decreases the supply of the CO_2_ needed for carbon assimilation via the Calvin-Benson cycle (CBC) and ultimately, starch synthesis (Lawlor et al., 2009). A limitation in carbon assimilation results in down-regulation of carbohydrate metabolism, which serves as an immediate precursor for the production of *e*.*g*. amino acids and/or energy donors such as nucleotides. Thus, the balancing of biochemical processes, especially carbohydrate and nitrogen metabolisms and the concomitant pathways including glycolysis and TCA cycle during stress is of great importance for plants to tolerate adverse conditions. Knowledge gained on the nature of plant stress responses allowed the development of various experimental strategies to improve drought tolerance (Pires et al., 2016; Fàbregas et al., 2018).

Limitations in the fixation of atmospheric CO_2_, whether caused by internal or external factors, will result in over-reduction of the photosynthetic electron transport chain (PETC) in chloroplasts, leading to inhibition of both PSI and PSII activities (Haupt-Herting and Fock, 2002). Once the availability of terminal electron acceptors becomes limiting, the PETC begins to leak electrons, resulting in the reduction of oxygen to detrimental compounds such as peroxides, superoxide and hydroxyl radicals, commonly classified as reactive oxygen species (ROS) (Takagi et al., 2016). The photorespiratory pathway of C3 plants represents a major sink for electrons under conditions of either limited CO_2_ availability or drought stress (Cruz de Carvalho, 2008). Also, the plastid terminal oxidase (PTOX) can extract electrons from plastoquinone (PQ), which are used to reduce oxygen to water, thereby maintaining the oxidation status of PSII during stress episodes (Sun and Wen, 2011).

To overcome the restriction of photosynthesis and thus the limitation of carbohydrate and metabolite production for better growth, we have been pursuing an alternative strategy by expressing specific cyanobacterial electron shuttles in chloroplasts (Tognetti et al., 2006; Zurbriggen et al., 2009). Among them, flavodiiron proteins (*Flvs*) represent a class of electron carriers able to reduce oxygen directly to water without ROS formation (Saraiva et al., 2004). Flavodiiron proteins have been found in many prokaryotic species (Wasserfallen et al., 1998) as well as in anaerobic protozoa, green algae and most plant lineages, with the major exception being angiosperms (Zhang et al., 2009; Peltier et al., 2010; Allahverdiyeva et al., 2015b).

In photosynthetic organisms, *Flvs* protect against photoinhibition by reducing oxygen in the non-heme diiron active site of their metallolactamase-like domain. The flavin mononucleotide (FMN) present in the C-terminal flavodoxin-like domain acts as a co-factor for this reaction, enabling electron transfer to the Fe-Fe centre (Silaghi-Dumitrescu et al., 2005). The genome of cyanobacterium *Synechocystis* sp. PCC 6803 (hereafter *Synechocystis*) encodes four distinct *Flv*s, *Flv1* through *Flv4* (Allahverdiyeva et al., 2011). *Flv1* and *Flv3* may form part of a single operon or be interspersed with 1 to 5 open reading frames (ORFs), whereas *Flv2* and *Flv4* are organized as an *Flv4*-ORF-*Flv2* operon. *Flv1* and *Flv3* have been proposed to form a heterodimer able to protect PSI under fluctuating light conditions by preventing the accumulation of ROS at the level of PSI (Helman et al., 2003; Allahverdiyeva et al., 2013, 2015a; Sétif et al., 2020). *Flvs* can mediate Mehler-like reactions and therefore complement cyclic electron transfer pathways in relieving the excess of excitation energy on the PETC (Dang et al., 2014; Gerotto et al., 2016), a phenomenon recently also observed in *Arabidopsis thaliana* plants expressing the *Flv1*/*Flv3* orthologues from the moss *Physcomitrella patens* (Yamamoto et al., 2016). When Gómez et al. (2018) introduced the *Synechocystis Flv1/Flv3* genes into tobacco, the proton motive force of dark-adapted leaves was enhanced, while the chloroplasts’ photosynthetic performance under steady-state illumination remained comparable to that of wild-type (WT) siblings. The heterologous expression of *P. patens Flv1* and *Flv3* in two rice mutants defective in cyclic electron transport was shown to restore biomass accumulation to WT levels (Wada et al., 2018). Recently, we demonstrated that the co-expression of *Synechocystis Flv1* and *Flv3* in *A. thaliana* enhanced the efficiency of light utilization, boosting the plant’s capacity to accumulate biomass as the growth light intensity was raised (Tula et al., 2020).

The present study aimed to create an additional dissipating electron sink downstream of PSI in the chloroplasts of barley, achieved by co-expressing *Synechocystis Flv1* and *Flv3*, and to determine the benefits that the presence of such transgenes could bring to the plant response to drought stress with respect to the production of carbohydrates and accompanying intermediates. Barley is the fourth most important cereal as a source for food and fodder and considered a model crop to investigate the influence of *Flv1* and *Flv3* expression on productivity traits such as biomass and yield. The focus was to investigate whether metabolic activity through photosynthesis can improve drought stress tolerance, thereby supporting the growth of plants exposed to this commonly occurring constraint over crop productivity.

## MATERIALS AND METHODS

### Barley transformation and growth

The methods used to transform barley followed those reported by Marthe et al. (2015). Briefly, the *Synechocystis Flv1* and *Flv3* genes were PCR-amplified, integrated into the pUBI-AB-M plasmid and subsequently cloned via the *Sfi*I restriction site into the binary vector p6i-2×35S-TE9 (Figure 1A), This generic vector harbours *hygromycin phosphotransferase* (*hpt*) as a plant selectable marker gene containing the potato *LS1* intron and driven by a double-enhanced Cauliflower Mosaic Virus (*CaMV*) *35S* promoter, the *Sm/Sp* (Streptomycin/Spectinomycin) bacterial selection marker gene and T-DNA borders derived from the p6i plasmid (DNA-Cloning-Service, Hamburg, Germany). Each *Flv* gene was placed between the maize *Polyubiquitin*-*1* promoter including 5’-untranslated region and first intron and the *Agrobacterium tumefaciens nos* terminator, with its coding region being fused in-frame at its 5’-end with a DNA fragment encoding the pea ferredoxin-NADP^+^ reductase (FNR) transit peptide for chloroplast targeting. The individual constructs harbouring either *Flv1* or *Flv3* were transformed into the barley cultivar ‘Golden Promise’ using *Agrobacterium tumefaciens* AGL-1 (a hypervirulent succinamopine strain with C58 background) by electroporation. Putative transgenic calli were kept for 12 h at 24°C in the light (mean relative humidity 50%) and for additional 12 h at 18°C in the dark (mean relative humidity 80%) until the formation of plantlets following shoot and root development. Thereafter, plantlets were transferred to soil and maintained at 80% humidity for 7 to 10 days by covering with a plastic hood. Plants were grown in a greenhouse providing a 12-h photoperiod at 250 µmol photons m^-2^ s^-1^ and a day/night temperature of 16°C/12°C (ambient conditions) until maturity, and grains were harvested for further experiments.

**FIGURE 1.**
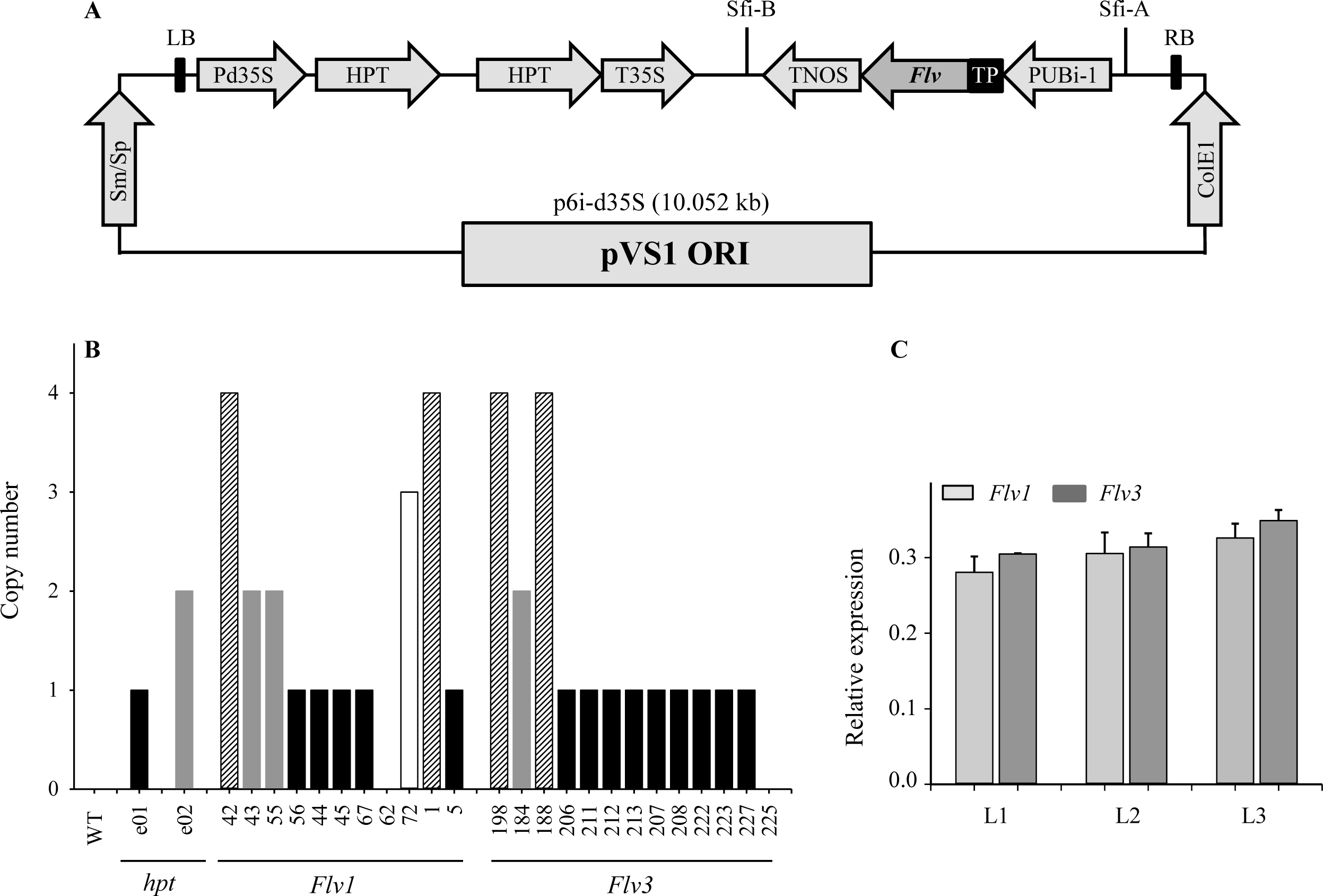
Expression of cyanobacterial *Flv1-Flv3* genes in barley plants. (A) Schematic representation of the p6i-2×35S-TE9 binary vector used to clone *Flv1* and *Flv3* genes. The vector harbours the *hpt* plant selectable marker gene containing the potato LS1 intron and driven by a double-enhanced *CaMV 35S* promoter (Pd35S) and terminator (T35) as well as the *Sm/Sp* bacterial selection marker gene. A sequence encoding the chloroplast-targeting FNR transit peptide (TP) was fused in-frame to the 5’-termini of *Flv1* and *Flv3* coding regions, and placed under control of the maize *Ubi*-*1* promoter with its 5’-untranslated regions and the first intron (PUbi-1) and the *nos* terminator between T-DNA borders derived from the p6i plasmid. (B) Determination of the copy number of individual *Flv* genes harboured by transgenic barley plants, as estimated using quantitative real-time PCR. T_1_ plants containing a single-locus of *Flv1* and *Flv3* (black bars) which could be identified by a 3:1 presence/absence PCR analysis were chosen as progenitors of the subsequently analyzed transgenic plants. WT barley represented the negative control and two plants (e01, e02) known to harbour one and two copies of the *hpt* gene, respectively, as positive controls. (C) Determination of *Flv* transcript levels in transgenic barley lines L1, L2 and L3 co-expressing *Flv1 and Flv3* genes. Data are shown in the form of means ± SE (*n* = 6).

T_1_ generation grains were sown in 96-well trays containing substrate 2 (Klasmann-Deilmann GmbH, Saterland, Germany), compost and sand (2:2:1), held at 4°C for 14 days, then exposed to a 16-h photoperiod at a day/night temperature of 18°C/12°C. Seedlings at the four-leaf stage were potted into a 3:2:1 compost, vermiculite and sand mixture and grown to maturity in a greenhouse under ambient conditions.

### Transgene copy number and *Flv* expression analysis

An estimate of the number of *Flv* transgene copies present in leaves of barley T_1_ individuals was obtained using a quantitative real-time PCR assay as described by Song et al. (2002) and Kovalchuk et al. (2013). Briefly, DNA was extracted from the second leaf of each plant following the method of Saghai-Maroof et al. (1984) and was serially diluted in sterile deionized water to give solutions containing between 12.5 and 200 ng µL^-1^ DNA. For the calculation of transgene copy number from unknown DNA samples, a serial dilution (400, 200, 100, 50 and 25 ng) of genomic DNA extracted from an available plant known to contain 1-2 copies of the *hpt* gene was used as the target sequence. Primers and PCR conditions are listed in Supplementary Table S1. For template loading normalization, the PCR reactions included a dual-labelled sequence 5′-CAL fluor Gold 540-ATGGTGGAAGGGCGGCTGTGABHQ1 as a probe complementary to a portion of the barley orthologue of the wheat *Pin-b* gene (Kovalchuk et al., 2013). The PCR efficiency for each primer set was determined from an analysis of the Ct values obtained from the serial dilution. Transgene copy numbers were determined by applying the 2^−ΔΔCT^ method (Li et al., 2004; Figure 1B). For each single-locus transgene construct harbouring either *Flv1* or *Flv3*, 16 T_1_ individuals were then self-pollinated. Homozygotes were selected by segregation analysis as determined by PCR amplification with primers *Flv1* F/R and *Flv3* F/R given in Supplementary Table S1. Only those behaving as having a single major gene (exhibiting a 3:1 segregation) in the T_2_ generation were retained as illustrated in Supplementary Figure S1. To monitor expression of the *Flv1/Flv3* genes in three independent lines (Figure 1C), total RNA was extracted from young leaves according to Logemann et al. (1987). RNA was subjected to DNase treatment (Thermo Fischer Scientific, Dreieich, Germany) and converted to single-stranded cDNA using a RevertAid first-strand cDNA synthesis kit (Life Technologies, Darmstadt, Germany) with a template of 1 µg total RNA and oligo (dt) primer. The reaction was run at 42°C for 60 min. Quantitative reverse transcription-PCR (qRT-PCR) was performed in a CFX384 touch real-time system (Bio-Rad, USA) using the SYBR Green Master Mix Kit (Bio-Rad, Feldkirchen, Germany). Primers employed to amplify *Flv1* (Flv1-RT F/R) and *Flv3* (Flv3-RT F/R), along with those amplifying the reference sequence gene *ubiquitin-conjugating enzyme 2* (E2 F/R) that was stably expressed under the experimental conditions tested for barley are listed in Supplementary Table S1. Relative transcript abundances were determined using the Schmittgen and Livak (2008) method. Each qRT-PCR result relied upon three biological replicates per line, each of which being represented by three technical replicates.

To produce double-homozygous plants harbouring *Flv1/Flv3*, single-locus T_2_ homozygotes with nearly same expression level were then inter-crossed following with two generations of self-pollination (Supplementary Figure S2). Siblings lacking *Flv* fragments, confirmed by PCR amplification (Figure S1), were used as ‘azygous’ control plants.

### Quantifying the barley response to drought stress

A representative set of barley plants harbouring *Flv1*/*Flv3* transgenes, were selected along with sibling azygous plants. A set of 24 plants of each of the *Flv1/Flv3* transgenic lines (F_3_), WT and azygous controls were grown for 28 days under a well-watered regime in a chamber providing ambient conditions. Twelve of the seedlings were then transferred into 5-cm pots with 50 g of soil (one seedling per pot) for the drought stress treatment at the vegetative stage and were allowed to recover for 3 days after being transferred. The other 12 seedlings were planted in larger pots (20-cm diameter and 200 g of soil, one seedling per pot) to assess the effect of stress at the reproductive stage. For the stress experiment at the seedling stage, six plants were kept under well-watered, ambient conditions, maintaining a soil moisture level of 65-70% of field capacity (FC) (Figure 2A). The remaining six plants were subjected to the drought treatment by withholding water for 3-4 days until the soil moisture level in the pots fall to 10-12% FC, and this state was maintained for five days (Figure 2B). Subsequently, the 12 treated plants were transferred to the glasshouse and grown under well-watered conditions until maturity (∼90 days) to determine growth parameters such as days to heading.

**FIGURE 2.**
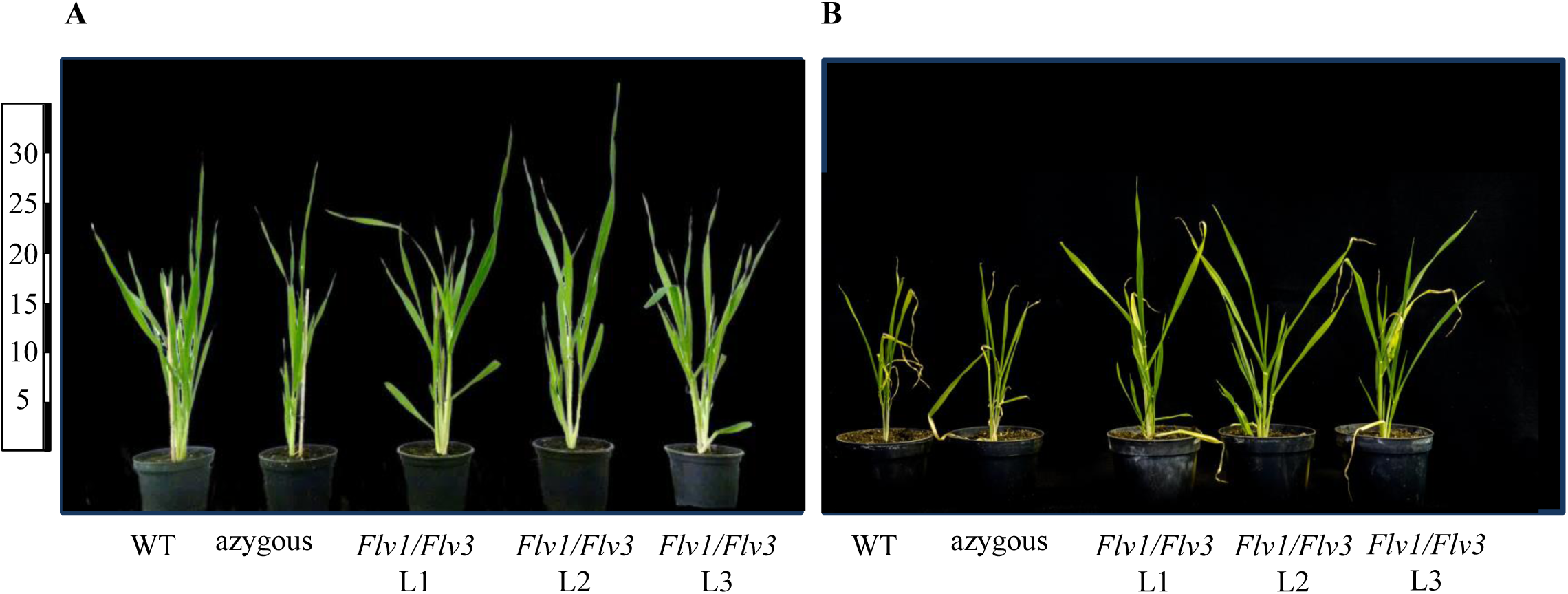
The appearance of typical barley plants heterologously expressing *Flv* genes at the seedling stage under ambient (A) and drought-stressed (B) conditions. Lines L1-L3 harbour both *Flv1* and *Flv3* genes. Images captured seven days after rewatering from a soil maintained at 10-12% FC for 5 days. Growth performance of WT, azygous and transgenic plants in ambient condition (A). Seven days after re-watering, WT barley plants exposed to severe drought exhibited retarded growth and leaf wilting, while leaves of the three transgenic lines retained turgor (albeit turning slightly yellowish). Numerals on the left indicate height in cm.

For the reproductive stage stress experiment, plants were kept well-watered (65-70% FC) under ambient conditions until the emergence of the first spike in 90% of the plants. Drought stress treatment was imposed five days post-anthesis by withholding water until FC fell to 10 to 12% and leaf wilting was observed. Thereafter, each pot was given 200 mL water every fourth day to maintain the soil moisture level at 10-12% FC over 21 days. Control plants were kept fully watered throughout. Flag leaves were collected 10 days after stress had been initiated, and the fresh weight (FW) of each leaf was measured immediately before it was placed into a collection tube. The relative water content (RWC) was calculated using 6 individuals each of WT and transgenic plants applying the following equation: RWC (%) = (FW − DW)/(TW − DW) × 100, where FW is the fresh weight at harvest time, TW is the total weight as total turgor estimated after 24 h of imbibition, and DW is the dry weight after 48 h at 85°C (Marchetti et al., 2019).

### Phenotypic effects of drought

The effect of drought stress on barley plants was assessed by measuring the following traits: days to heading, defined as the number of days from sowing to the time when 50% of the spikes had emerged from the flag leaf sheath, using Zadoks scale 55 (Zadoks et al., 1974); plant height (the height from the soil surface to the tip of the longest spike, excluding awns); above-ground plant biomass at maturity measured after the plants had been oven-dried at 60°C for 72 h); the number of spikes produced per plant; the grain number per plant and the grain yield (the weight of total grains per plant). The latter two traits were quantified using a Marvin-universal seed analyser (GTA Sensorik GmbH, Neubrandenburg, Germany).

### Metabolite measurements

Due to the importance of the flag leaf in grain filling compared to other leaves in barley (Shahinnia et al., 2019), flag leaves of two spikes per plants with the same developmental stage were sampled for metabolite determinations when a completed leaf rolling as the primary visible symptom of drought stress occurred. The contents of individual amino acids, including the stress marker proline, were quantified as described by Mayta et al. (2018), whereas extraction and analysis of soluble sugars were essentially performed according to Ahkami et al. (2013).

Adenine nucleotides were quantified employing an UPLC-based method developed from that described by Haink and Deussen (2003). Prior to the separation step, a 50-µL aliquot of the sample and a mixture of ATP, ADP and AMP were derivatized by the addition of 25 µL of 10% (v/v) chloracetaldehyde and 425 µL of 62 mM sodium citrate/76 mM KH_2_PO_4_, pH 5.2, followed by a 40-min incubation at 80°C, cooling on ice, and centrifugation at 20,000 *g* for 1 minute. The separation was achieved using an ultra-pressure reversed-phase chromatography system (AcQuity H-Class, Waters GmbH, Eschborn, Germany) consisting of a quaternary solvent manager, a sample manager-FTN, a column manager and a fluorescent detector (PDA eλ Detector). The gradient was established using eluents A (TBAS/KH_2_PO_4_: 5.7 mM tetrabutylammonium bisulfate/30.5 mM KH_2_PO_4_, pH 5.8) and B (a 2:1 mixture of acetonitrile and TBAS/KH_2_PO_4_); the Roti C Solv HPLC reagents were purchased from Roth (Karlsruhe, Germany). The 1.8 µm, 2.1×50 mm separation column was a Luna Omega C18, (Phenomenex, Aschaffenburg, Germany). The column was pre-equilibrated for at least 30 minutes in a 9:1 mixture of eluents A and B. During the first two minutes of the run, the column contained 9:1 A:B, changed thereafter to 2:3 A:B for 2 minutes followed by a change to 1:9 A:B for 1 minute and set to initial values of 9:1 for 2 minutes. The flow rate was 0.5 mL min^-1^ and the column temperature was maintained at 45°C. The excitation and emission wavelengths were 280 nm and 410 nm, respectively. Chromatograms were integrated using Empower Pro software (Waters, Eschborn, Germany). Energy charge was calculated from the expression ([ATP] + 0.5 [ADP])/([ATP] + [ADP] + [AMP]) (Atkinson, 1967).

### Statistical analyses

Descriptive statistics (means and SE) and data analysis were carried out using SigmaPlot (Systat Software, San Jose, CA, USA). The Student’s *t*-test was applied for evaluating statitically significant differences between means of individual transgenic lines versus the wild-type.

## RESULTS

### *Flv* transgenes improved the productivity of barley plants subjected to drought stress at the seedling stage

When grown under ambient conditions, *Flv*-expressing plants were taller than their WT siblings (Figures 2A and 3A), without significant differences in aboveground biomass dry weight (Figure 3B). Height differences between WT and transgenic plants were maintained under drought stress applied at the seedling stage (Figure 2B and 3A). The treatment caused a major decrease (up to 40%) of total biomass in non-transformed and azygous plants, which was reduced to less than 10% in their transgenic siblings (Figure 3B). Compared to WT plants, up to 1.5-fold more biomass was accumulated by *Flv1/Flv3*-expressing lines under drought (Figure 3B). In the absence of stress, *Flv1*/*Flv3* transgenic plants generally reached heading 2-3 days sooner than non-transformed and azygous counterparts, with these differences becoming more pronounced (5-7 days) under drought (Figure 3C).

**FIGURE 3.**
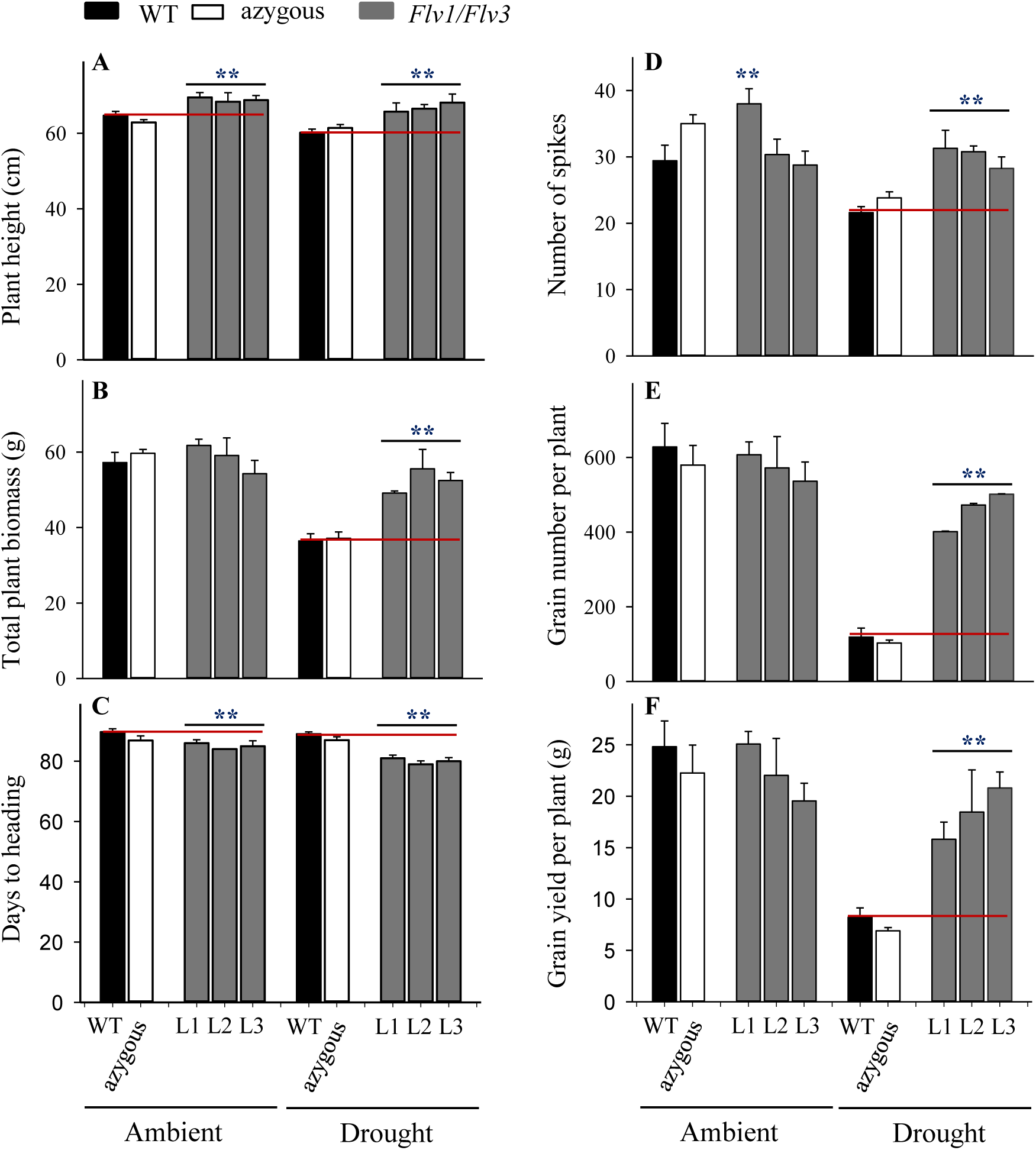
Effect of heterologously expressing *Flv1/Flv3* genes on productivity-associated traits of barley plants grown either under ambient conditions or exposed to drought stress for 5 days at the seedling stage. Measurements were carried out at maturity (∼90 days). Other experimental details are given in Materials and Methods. (A) Plant height, (B) total plant biomass, (C) days to heading, (D) number of spikes per plant, (E) total number of grains per plant, (F) overall grain yield per plant. Lines L1-L3 co-express *Flv1 and Flv3* genes. Data are shown as means ± SE (*n =* 6). **: means differed significantly (P ≤ 0.01) from those of non-transgenic plants.

Plants expressing both transgenes were the least compromised by drought stress with respect to the number of spikes produced (Figure 3D). Compared to WT and azygous plants, there was also a significant preservation in the number of grains set per plant by drought-challenged *Flv1/Flv3* transgenic lines. The stress treatment decreased grain number by as much as 4-fold in WT and azygous plants while the three transgenic lines displayed less than 20% reduction (Figure 3E), setting at least 3.7-fold more grain than their non-transgenic controls in drought-stressed conditions (Figure 3E). A similar trend was observed for total grain yield, which was reduced up to 3-fold in WT and azygous plants upon drought stress, but only up to 30% in the transformants (Figure 3F). Indeed, the grain yield of *Flv1/Flv3* transgenic plants from lines L2 and L3 appeared not to be affected by the adverse condition. Total grain yield per plant was up to 3-fold higher in the *Flv1/Flv3*-expressing lines subjected to drought stress than that achieved by the non-transgenic plants (Figure 3F).

### *Flv* transgenes prevented yield loss in barley exposed to drought stress at the reproductive stage

The increased height of the *Flv1/Flv3* transgenic plants under non-stressed conditions was maintained as plants entered the reproductive stage (Figure 4A). While the relative water content measured at this stage decreased upon drought stress, it did not differ significantly between WT and transgenic plants grown under ambient conditions (about 78%) nor in plants exposed to drought stress (about 47%). The height increase driven by *Flv1/Flv3* presence was lost upon drought exposure at the reproductive stage (Figure 4A). In contrast, drought-dependent reduction in aboveground biomass was similar to that observed upon stress application at the seedling stage and was equally protected by *Flv1/Flv3* (Figure 4B). The imposition of drought stress at the reproductive stage advanced heading only in line L3 of *Flv* transgenic plants by around three days (Figure 4C).

**FIGURE 4.**
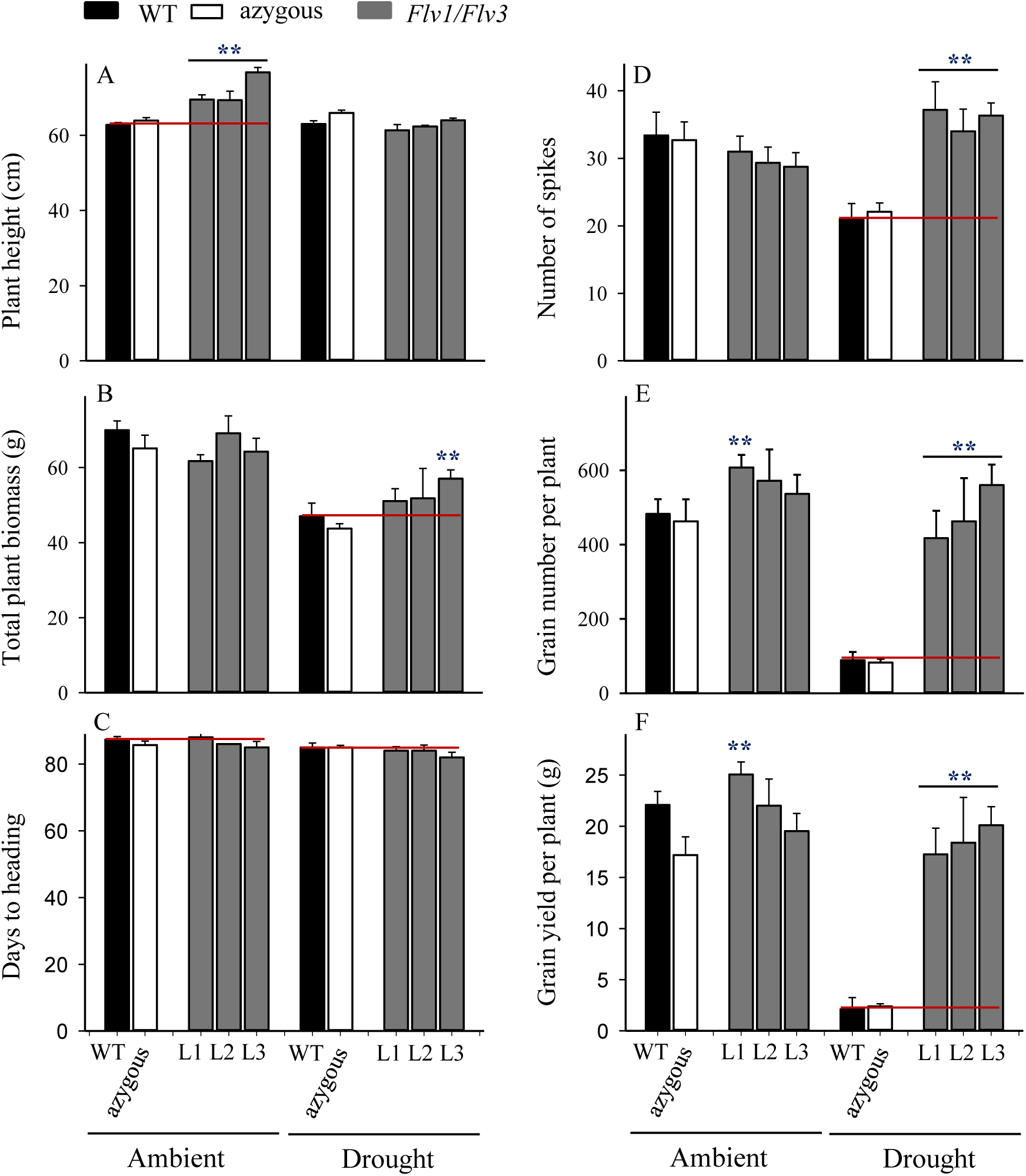
Effect of heterologously expressing *Flv1/Flv3* genes on productivity-associated traits of barley plants grown either under ambient conditions or exposed to drought stress for 21 days at the reproductive stage. Measurements were carried out at the end of the 21-day drought treatment. Other experimental details are given in Materials and Methods. (A) Plant height, (B) total plant biomass, (C) days to heading, (D) number of spikes per plant, (E) total number of grains per plant, (F) overall grain yield per plant. Lines L1-L3 harbour both *Flv1 and Flv3* genes. Data are shown as means ± SE (*n =* 6). **: means differed significantly (P ≤ 0.01) from those of non-transgenic plants.

With respect to the number of spikes produced per plant, the *Flv1/Flv3* transgenic plants were notable for the protective effect exerted under drought, while there was no variation between lines in the absence of stress (Figure 4D). Drought also had a devastating effect on yield when applied at the reproductive stage, but *Flv1/Flv3* transgenic plants were able to set ∼2-3-fold more grain per plant than their WT siblings (Figure 4E), and their grain yield was 8-to 9.5-fold greater (Figure 4F). Under these conditions, the grain yields of lines L2 and L3 were actually unaffected by the stress treatment. In summary, expression of *Flv1/Flv3* preserved major productivity traits such as the number of spikes, grain number and grain yield per plant in transgenic barley plants exposed to drought treatments applied at either the seedling or the reproductive stages (Figures 3 and 4).

### The effect of expressing *Flv* transgenes on carbohydrate contents and amino acid levels of drought-stressed barley plants

Under ambient conditions, flag leaf glucose and fructose were not detectable in control plants used for drought stress experiments applied at seedling stage or in a low amount at the reproductive stage, with no significant differences between WT, azygous and transgenic plants (Figure 5A, B, D and E). Sucrose also failed to display differences between lines, although their levels increased ∼5-fold as the plants challenged at the reproductive stage (Figure 5C and F).

**FIGURE 5.**
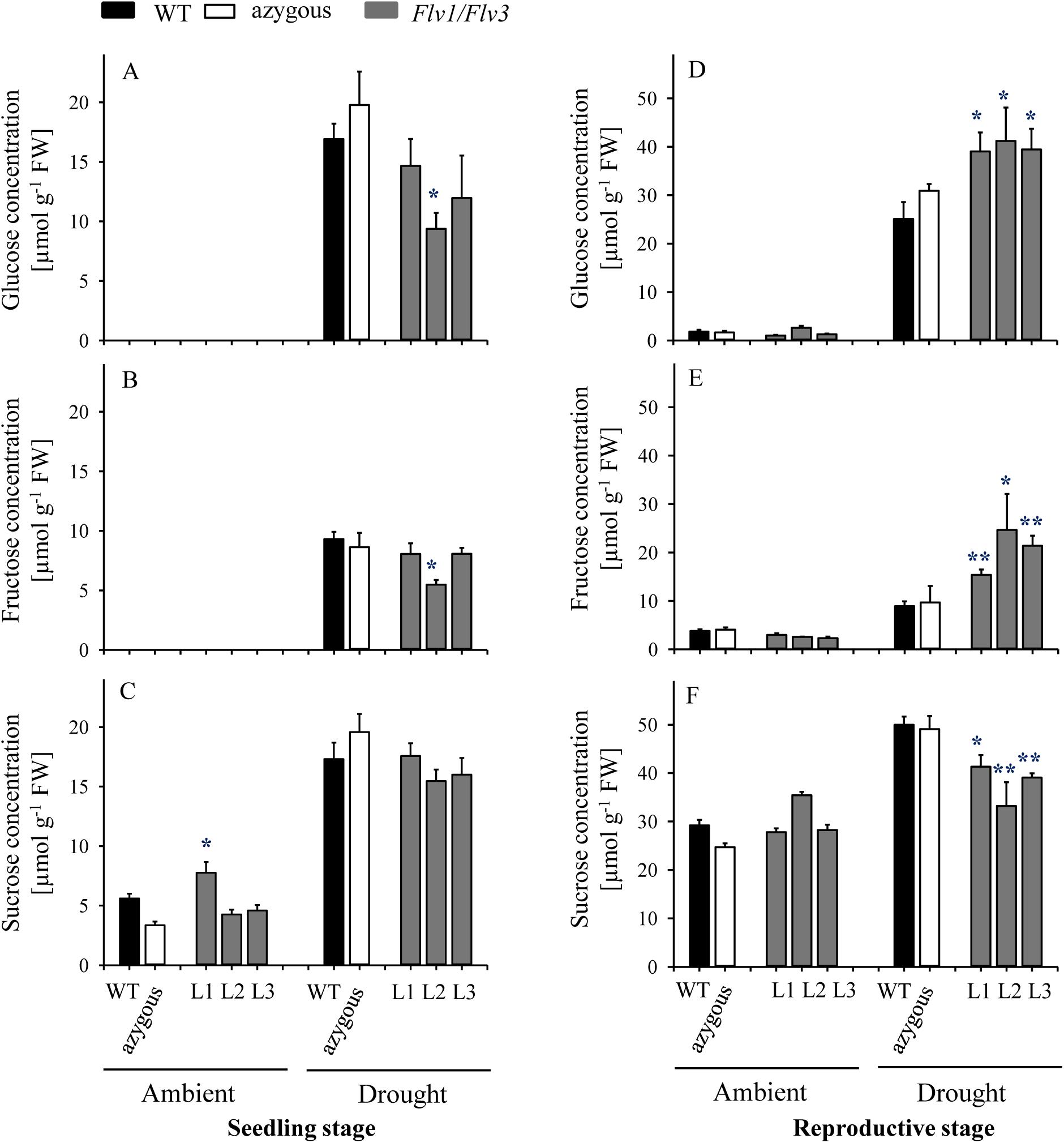
Effect of heterologously expressing *Flv1/Flv3* genes on sugar contents in flag leaves of barley plants grown either under ambient conditions or exposed to drought stress at the seedling stage (A-C) and the reproductive stage (D-F). Samples were collected at the leaf rolling stage. Other details are given in Materials and Methods. (A, D) Glucose, (B, E) fructose, (C, F) sucrose. Lines L1-L3 co-express *Flv1 and Flv3* genes. Data are shown as means ± SE (*n =* 6). **; *: means differed significantly (P ≤ 0.01 and P ≤ 0.05, respectively) from those of non-transgenic plants. FW, fresh weight.

Application of the drought treatment at the seedling stage led to major increases in all soluble sugars, irrespective of the genotype (Figure 5A-C). Significant differences between lines became instead apparent when the stress treatment was assayed at the reproductive stage, with higher leaf glucose and fructose contents (Figure 5D and E) and lower sucrose levels in transgenic plants compared to their WT siblings (Figure 5F).

In plants of drought stress applied at the seedling stage, flag leaf amino acid contents were not affected by *Flv1/Flv3* expression when plants had been grown under ambient conditions except for the case of glutamate, whose levels were up to 1.6-fold higher relative to WT counterparts (Figure 6A; Supplementary Table S2). Drought treatment had little effect on the amounts of free amino acids in WT and azygous plants, but for a 4-fold increase in glycine (Figure 6A-E). In contrast, an increased pool of histidine, asparagine, serine, glutamine, glutamate, asparagine, threonine and alanine was observed in *Flv1/Flv3* transgenics under stress conditions (Figure 6A-E; Supplementary Table S2). Leaf contents of proline increased strongly (up to 60-fold) in drought-exposed WT and azygous plants, which is in line with its recognized role as a stress marker. By contrast, proline levels increased significantly less in stress-treated *Flv* transformants, despite their higher proline levels under ambient conditions (Figure 6F, inset).

**FIGURE 6.**
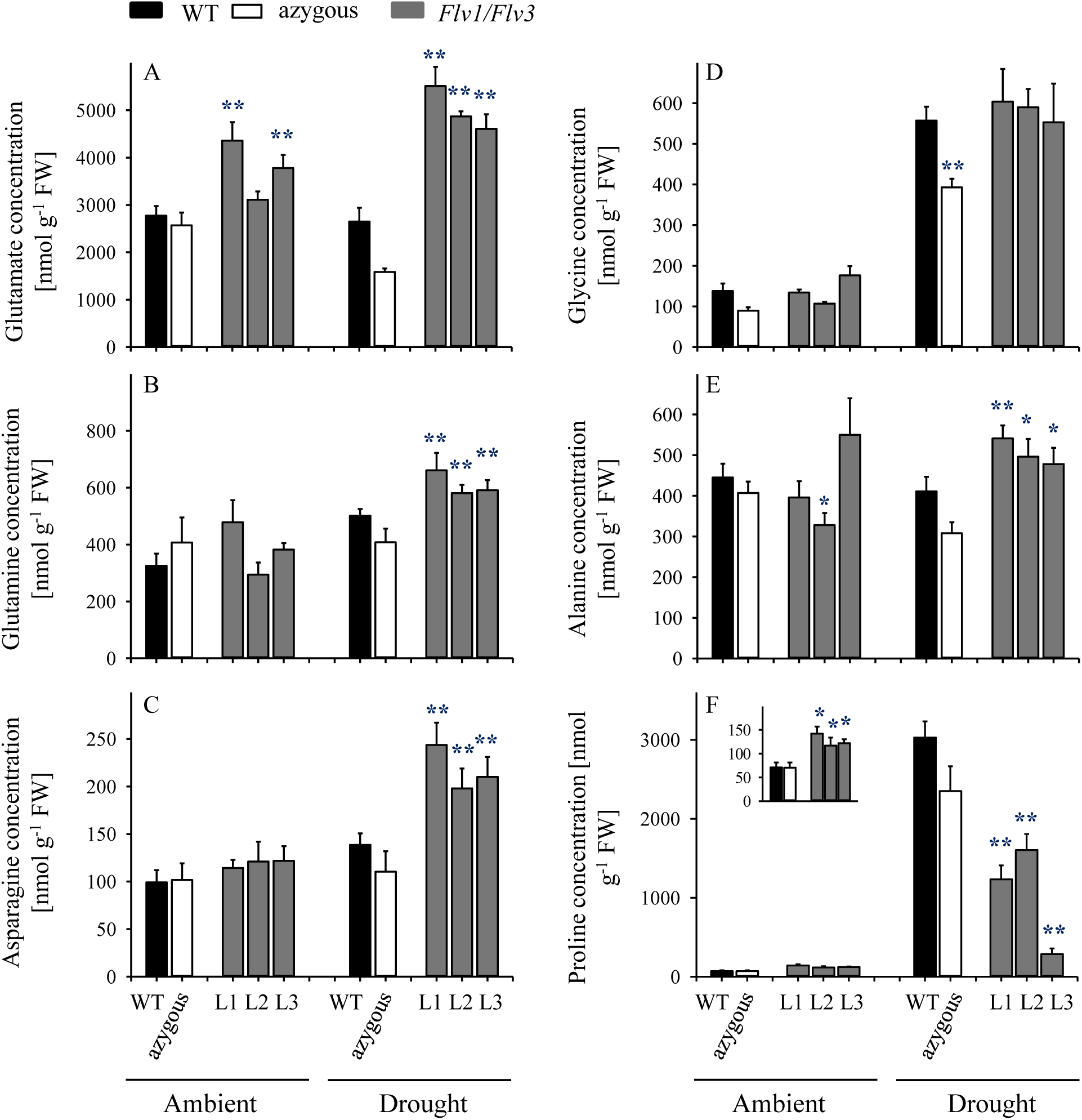
Effect of heterologously expressing *Flv1/Flv3* genes on free amino acid contents in flag leaves of barley plants grown either under ambient conditions or exposed to drought stress **at the seedling stage**. Amino acid levels were measured in the same samples used for carbohydrate determinations. (A) Glutamate, (B) glutamine, (C) asparagine, (D) glycine, (E) alanine and (F) proline. Lines L1-L3 harbour both *Flv1 and Flv3* genes. Data are shown as means ± SE (*n = 5-*7). **; *: means differed significantly (P ≤ 0.01 and P ≤ 0.05, respectively) from those of non-transgenic plants. FW, fresh weight.

Under ambient conditions, the flag leaf contents of free amino acids increased significantly as the plants entered the reproductive stage (Figure 7; Supplementary Table S3), with no major differences between lines except for proline and glutamine, which accumulated to lower levels in *Flv*-expressing plants (Figure 7B and F). Drought exposure increased the amounts of several amino acids (most conspicuously proline) in WT and azygous plants, (Figure 7; Supplementary Table S3). Noteworthy, the stress condition did not affect the amounts of specific amino acids derived from the glycolytic metabolism, such as glutamate, glutamine, asparagine, aspartate and serine (Figure 7A-E), as well as glycine and threonine (Supplementary Table S3) in leaves of the *Flv* transformants. Proline levels were up-regulated by drought, but significantly less than in WT and azygous plants (Figure 7F). No clear differences were observed for other amino acids following exposure to drought as compared to non-stressed plants (Supplementary Table S3).

**FIGURE 7.**
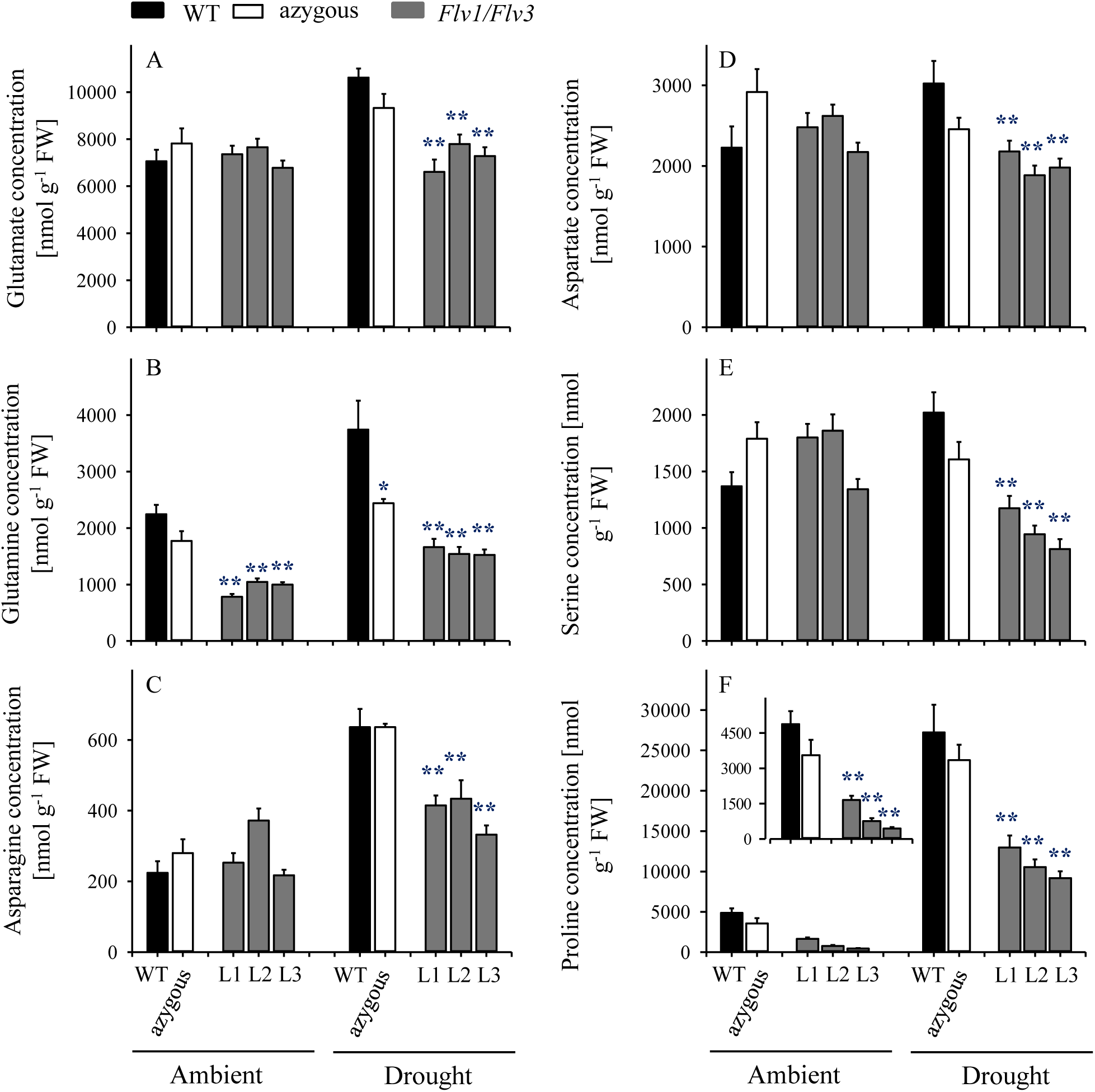
Influence of heterologously expressing *Flv1/Flv3* genes on free amino acid contents in flag leaves of barley plants grown either under ambient conditions or exposed to drought stress **at the reproductive stage**. Amino acid levels were measured in the same samples used for carbohydrate determinations. (A) Glutamate, (B) glutamine, (C) asparagine, (D) aspartate, (E) serine and (F) proline. Lines L1-L3 co-express *Flv1 and Flv3* genes. Data are shown as means ± SE (*n = 6-*7 for WT and azygous, and *n* = 8-14 for transgenic lines). **; *: means differed significantly (P ≤ 0.01 and P ≤ 0.05, respectively) from those of non-transgenic plants. FW, fresh weight.

### The effect of expressing *Flv* transgenes on the energy status of drought-stressed barley plants

At the seedling stage, ATP and ADP contents were similar in leaves from WT, azygous and transgenic plants under ambient conditions while there was a decrease of AMP levels up to 1.7-fold in *Flv*-expressing lines compared to WT and azygous siblings (Supplementary Figure S3A-C). The contents of all adenylates strongly increased in drought-stressed WT and azygous plants and in the transgenic line L1, whereas lines L2 and L3 maintained ATP and ADP at WT levels (Supplementary Figure S3A-C).

Upon reaching the reproductive stage, adenylate contents increased 3-to 8-fold in WT and azygous plants under ambient conditions, but significantly less in the transformants (Supplementary Figure S3D-F). Accordingly, adenine nucleotide levels were as much as 3-fold (AMP), 1.8-fold (ADP) and 2.1-fold (ATP) lower in the leaves of *Flv*-expressing plants compared to WT and azygous counterparts (Supplementary Figure S3D-F). Drought stress, in turn, led to a moderate decline in adenylate contents (especially ADP and AMP) in WT and azygous plants but increased those of *Flv* transformants, resulting in similar levels for the three nucleotides in all lines (Supplementary Figure S3D-F).

As a consequence of these effects of *Flv1*/*Flv3* expression on adenylate levels, the ATP/ADP ratio and the energy charge were largely similar between lines under both ambient and drought conditions applied at either the seedling or reproductive stages, with only few exceptions illustrated in Supplementary Figure S4.

## DISCUSSION

This is the first study to show that introduction of the cyanobacterial *Flv1 and Flv3* genes into the chloroplast improves the productivity of barley under drought through maintenance of metabolic activity and increasing carbohydrate and amino acid utilization.

### The heterologous expression of *Flv1/Flv3* in barley improves plant productivity under drought stress

Crops frequently encounter drought as transient or terminal stress (Alegre, 2004). Plant survival under these unfavourable conditions depends on their duration and intensity. When exposed to moderate stress, plants survive by adaptation or acclimation strategies and by repair mechanisms. To cope with chronic drought conditions causing severe damage or death, they evolve resistance mechanisms further classified into drought avoidance and drought tolerance (Price et al., 2002). A typical response of cereals such as barley to drought or high-temperature stress is to slow down their vegetative growth, followed by progressive leaf wilting if the adverse condition is prolonged. When these stresses occur around anthesis, the plant response may include premature leaf senescence, which results in a decline in photosynthesis and assimilate production as well as an acceleration of physiological maturation (Gan, 2003). Under terminal drought, crop yields are limited by a combination of infertility, grain abortion and reduced grain size (Sreenivasulu et al., 2007). Here, when barley plants were exposed to drought at the seedling stage, the heterologous expression of *Flv1*/*Flv3* resulted in the acceleration of heading time and flowering (Figure 3C). For such plants, one likely consequence is that they are less prone to experience terminal drought stress because they earlier reach maturity. The presence of the *Flv1/Flv3* transgenes was thus associated with the production of more spikes and a significantly higher grain number and yield under drought stress conditions applied at both the seedling and reproductive stages (Figures 3 and 4).

The combination of Flv1 and Flv3 proteins has been reported as being necessary to provide an effective electron sink under adverse environmental conditions in cyanobacteria. Allahverdiyeva et al. (2015b) have shown that cyanobacterial *Flvs* can act as heterodimers to facilitate a more rapid transfer of electrons to oxygen under conditions of excessive light. Loss-of-function mutants for both *Flv1* and *Flv3* in *Synechocystis* sp. PCC 6803 and *Anabaena* sp. PCC 7120 are compromised in their growth and in their ability to photosynthesize when exposed to fluctuating light (Allahverdiyeva et al., 2015a). It was proposed that this behaviour is related to a malfunction of PSI, which induces ROS production and hence causes oxidative stress (Allahverdiyeva et al., 2015a). However, an alternative scenario is that the key consequence resulting from *Flv* deficiency is a reduction in ATP abundance derived from photosynthesis, as evidenced by the effect of low light intensities on the energization of the membrane (Allahverdiyeva et al., 2013). Under conditions of drought stress, the barley *Flv1/Flv3* transgenic plants out-performed their non-transgenic controls in the accumulation of aboveground biomass, the number of grains set and the grain yield per plant (Figures 3 and 4). These observations suggest that heterodimeric Flvs are also functional in a monocotyledonous species, acting to maintain growth in a situation where surplus electrons are produced. Additional support for this contention is also provided by the reduced accumulation of proline (a marker of drought stress, see Szabados and Savouré, 2010) in leaves of the transgenic plants (Figures 6 and 7).

### The heterologous expression of *Flv1/Flv3* resulted in a distinct response of carbohydrates, amino acids and energy status at various developmental stages in drought-stressed barley plants

Drought stress suppresses the production of carbohydrates either by restricting CO_2_ fixation following to stomatal closure (Quick et al., 1992; Brestic et al., 1995), or via limiting the supply of ATP as a result of inhibition of ATP synthase (Tezara et al., 1999). Sucrose synthesized during photosynthesis represents the major feedstock for starch production (Counce and Gravois, 2006), but in drought-stressed plants it also acts as an osmolyte, helping to maintain turgor pressure and to mitigate membrane damage (Couée et al., 2006). The response of plants with respect to sugar accumulation under drought conditions depends on the species and even on the intraspecific lines within a given species, as reported for wheat by Guo *et al*. (2018). The comparison of drought-sensitive and -tolerant wheat varieties revealed that soluble sugars such as sucrose or fructose displayed opposite stress behaviour, that is, they are reduced in the sensitive and increased in the tolerant plants under drought (Guo et al., 2018). In the present study, drought treatments applied at either the vegetative or reproductive stages resulted in a strong accumulation of soluble sugars including glucose, fructose and sucrose in WT and transgenic plants (Figure 5). This indicates that these metabolites play important roles in the delivery of assimilates to sink organs for further growth (Fàbregas and Fernie, 2019, and references therein) or as osmo-protectants (Singh et al., 2015), and as such are highly sensitive markers of environmental adversities. Sugar accumulation is a general response to drought stress in different plant species, as demonstrated in the current study and several other reports (Singh et al., 2015; Das et al., 2017; Fàbregas et al., 2018; Fàbregas and Fernie, 2019). Remarkably, transgenic lines expressing *Flv1/Flv3* genes exhibited even higher glucose and fructose contents and a slightly lower sucrose content compared to those of WT and azygous plants under drought conditions (Figure 5), suggesting a higher activity of downstream pathways including glycolysis to keep pace with the environmental changes.

Improved metabolic activity exerted by chloroplast-expressed *Flv1/Flv3* is also reflected by the differential drought response of amino acid turnover. A schematic model describing metabolic fluxes in WT and Flv1/Flv3-transgenic plants is shown in Figure 8. At the vegetative stage, several amino acids such as Glu, Gln, Asp and Ala increased in the flag leaves of transgenic plants under drought with respect to those in WT siblings. By contrast, at the reproductive stage, most amino acids including Glu, Gln, Ser, Asp and Asn decreased while being maintained at the levels found in the absence of stress (Figure 6 and 7; Supplementary Tables S2 and S3). This contrasting effect of drought on amino acid accumulation (Figure 8) might be due to the fact that at the vegetative stage, barley plants invest all the assimilates into the defense mechanisms to resist the stress condition for better growth. Improved assimilate production in transgenic plants might result from a better performance of photosynthetic activity exerted by the presence of Flv1/Flv3 proteins as demonstrated in several studies (Yamamoto et al., 2016; Gómez et al., 2018; Wada et al., 2018).

**FIGURE 8.**
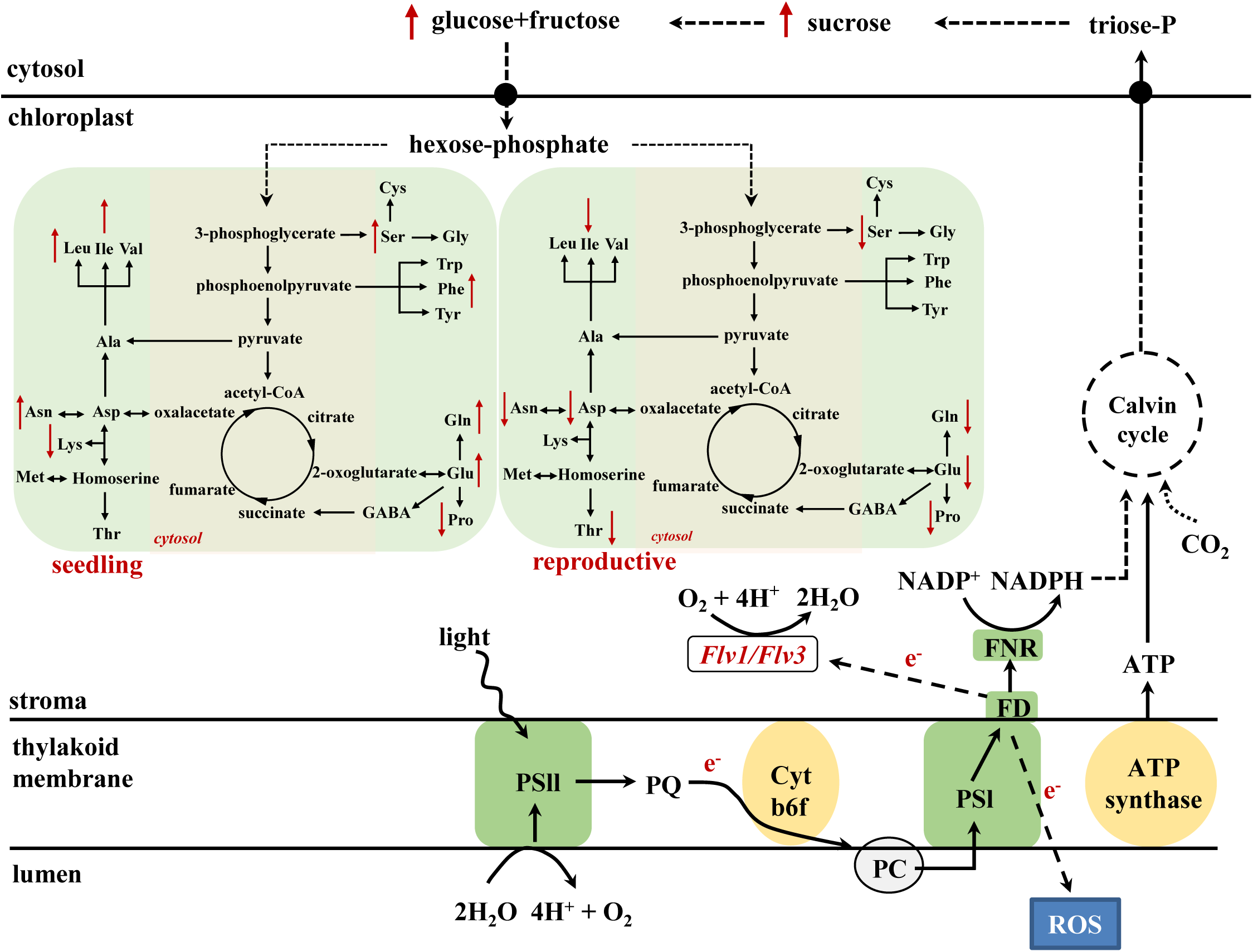
A model describing the metabolic consequences of heterologously expressing *Flv1/Flv3* genes in the chloroplasts of barley plants exposed to drought stress at the seedling (left) and at the reproductive stage (right). The presence of *Flv* gene products generates an electron sink and balances the electron pressure generated under stress by delivering the surplus of reducing equivalents to oxygen, which is converted to water. Based on the results, we propose that this activity is acting as a valve to relieve the excess of electrons and does not affect NADPH production, allowing CO_2_ assimilation through the Calvin-Benson cycle to form triose-phosphates. Sucrose produced from triose-phosphates is cleaved to soluble sugars glucose and fructose. The resulting hexose-phosphates are incorporated into amino acids, or used for energy production through glycolysis. As a consequence, these intermediates are preferentially employed to maintain the energy source necessary to support the growth of plants exposed to stress. PQ, plastoquinone; Cytb6/f, cytochrome *b*_6_/*f* complex; PC, plastocyanin; FD, ferredoxin; FNR, ferredoxin-NADP^+^ reductase.

At the reproductive stage, water limitation led to a strong increase in amino acid levels in WT flag leaves compared to non-stressed conditions (Figures 6 and 8; Supplementary Table S2). However, in the flag leaves of transgenic plants, the same amino acids were maintained at the levels found under non-stressed conditions or decreased in comparison to the contents of WT plants (Figures 7 and 8; Supplementary Table S3). At this stage, a stable metabolic activity is crucial for the maintenance of assimilate translocation from the flag leaves to the growing sink tissues, in this particular case the grains that are highly dependent on the delivery of the assimilates from the source organs. Thus, most likely WT plants use the produced sugars to synthesize amino acids such as Glu that serve as a key hub for the production of defense compounds such as proline, a sensitive marker of drought stress (Fàbregas and Fernie, 2019). However, due to a better performance of metabolic activity, transgenic barley plants may compensate the loss of nitrogen-containing amino acids by reducing proline production (Figure 6 and 7) and by using this saving for further assimilation and translocation to sink organs (Rai and Sharma, 1991; Hildebrandt et al., 2015).

Recent publications have demonstrated that high levels of energy and sugars improve plant development and tolerance to drought stress (Guo et al., 2018; Fàbregas and Fernie, 2019). This is also a fundamental basis for an active metabolism with increased pools of intermediates such as amino acids. Furthermore, amino acids have been reported to contribute to both membrane permeability and ion transport in the leaves of *Vicia faba* (Rai and Sharma, 1991), and to provide a source of energy (Hildebrandt et al., 2015). In particular, proline accumulation is usually induced during different environmental stresses, as it serves both as osmolyte and antioxidant (Szabados and Savouré, 2010). Here, the *Flv1/Flv3* transgenic barley plants accumulated less proline in their leaves than their WT controls when exposed to drought (Figure 6F and 7F), suggesting that they were capable of coping better with the stress than their WT counterparts. Moreover, *Flv*-expressing plants showed significant drought-associated increases in specific amino acids such as alanine, glutamate, serine and aspartate (Figure 8) which are derived from precursors of the glycolytic metabolism and serve as immediate primary substrates to build up nitrogen sources like glutamine and asparagine or antioxidative compounds like glutathione or polyamines. Thus, the metabolite profiling supports the idea that carbohydrates and amino acid metabolism help maintain the fitness of plants under drought stress, which is also in agreement with previously reported results of drought-tolerant varieties in other species (Guo et al., 2018).

Following exposure to drought, ATP levels were found to increase (relative to ambient conditions) in the leaves of *Flv* transgenic plants at both the seedling and reproductive stages (Supplementary Figure S3), indicating that *Flv1/Flv3* were able to maintain linear electron flow and thereby support ATP synthesis under the adverse condition. Sustaining cellular metabolism and ensuring growth and survival under stress rely heavily on a continuous supply of ATP (Sharkey et al., 1982). By improving the availability of electron acceptors at PSI, *Flv1/Flv3* can prevent ROS build-up (Rutherford et al., 2012), which may in turn inhibit both PSI and PSII activity and compromise the function of the ATP synthase complex (Lawlor, 1995).

## Conclusion

Data presented here show how integrating additional electron sinks to the PETC can boost the level of drought tolerance in a monocotyledonous crop species, irrespective of whether the drought condition was applied at the seedling stage or post-flowering. The heterologous expression of both *Flv1* and *Flv3* in barley had the effect of allowing efficient utilization of produced assimilates including sugars and amino acids, thereby supporting plant growth in the face of either early or late-onset drought and ultimately supporting the conversion of assimilates into biomass and yield (Figure 8). Overall, the experiments have confirmed that adopting this genetic manipulation approach has substantial potential to enhance the level of stress tolerance exerted by crop plants.

## DATA AVAILABILITY STATEMENT

This article does not contain any studies with human participants or animals performed by any of the authors. All datasets generated for this study are included in the article/Supplementary Material.

## CONFLICT OF INTEREST

The authors declare that they have no conflicts of interest.

## AUTHOR CONTRIBUTIONS

FS and MRH have made substantial contributions to conception and design, interpretation of the results and preparation of the manuscript. FS conducted the experiments and analysed the data. ST, GH, JK supported producing of transgenic plants. NR and NN helped in the phenotypic evaluation of the plants. RG, AFL and NC have been involved in the interpretation of the data and editing the manuscript. NC reviewed the manuscript. All authors read and approved the final manuscript for publication.

## FUNDING

The project was supported by funds of the Bundesministerium für Bildung und Forschung (BMBF), Germany, to FS, ST and MRH.

## ACKNOWLEDGMENTS

We wish to thank Melanie Ruff, Nicole Schäfer, Sabine Sommerfeld, Heike Büchner and Heike Nierig for their excellent technical assistances at the IPK. RG was a Fellow and AFL and NC are Staff Researchers from CONICET, Argentina. AFL and NC are Faculty members of the Facultad de Ciencias Bioquímicas y Farmacéuticas, Universidad Nacional de Rosario, Argentina.

## SUPPLEMENTARY FIGURES

**FIGURE S1.**
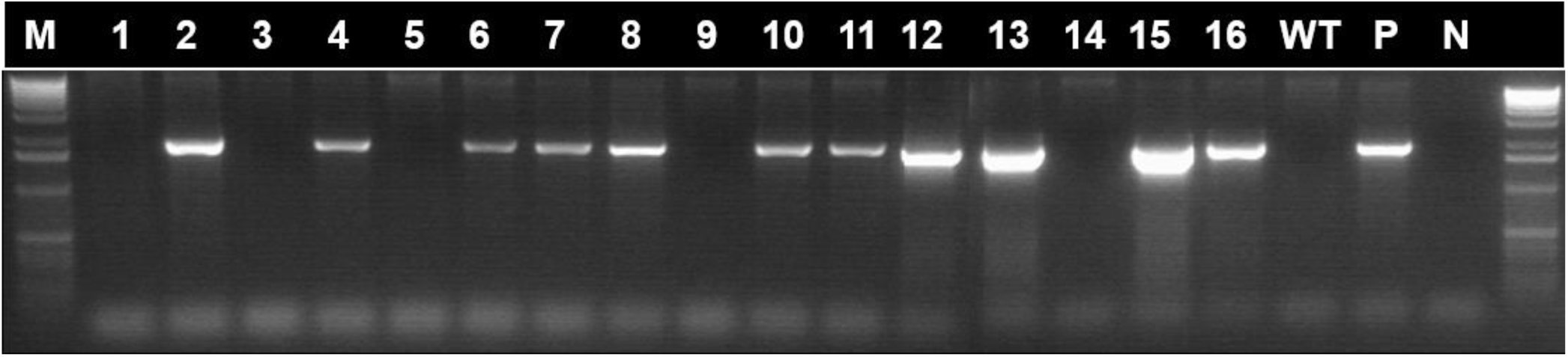
A representative segregation analysis of the *Flv1* transgene in a set of barley T_2_ individuals (lanes 1-16), as determined by PCR amplification. The selection of single-locus transgenic plants was made based on a monogenic (3:1) ratio for both *Flv1* and *Flv3*. M: 1 kbp DNA ladder, WT: wild-type, P: empty plasmid control, N: no-template negative control. The size of the target amplicon was 1.8 kbp. Plants lacking the *Flv1* amplicon (*i*.*e*. lanes 1, 3, 5, 9, 14) were used to produce azygous individuals.

**FIGURE S2.**
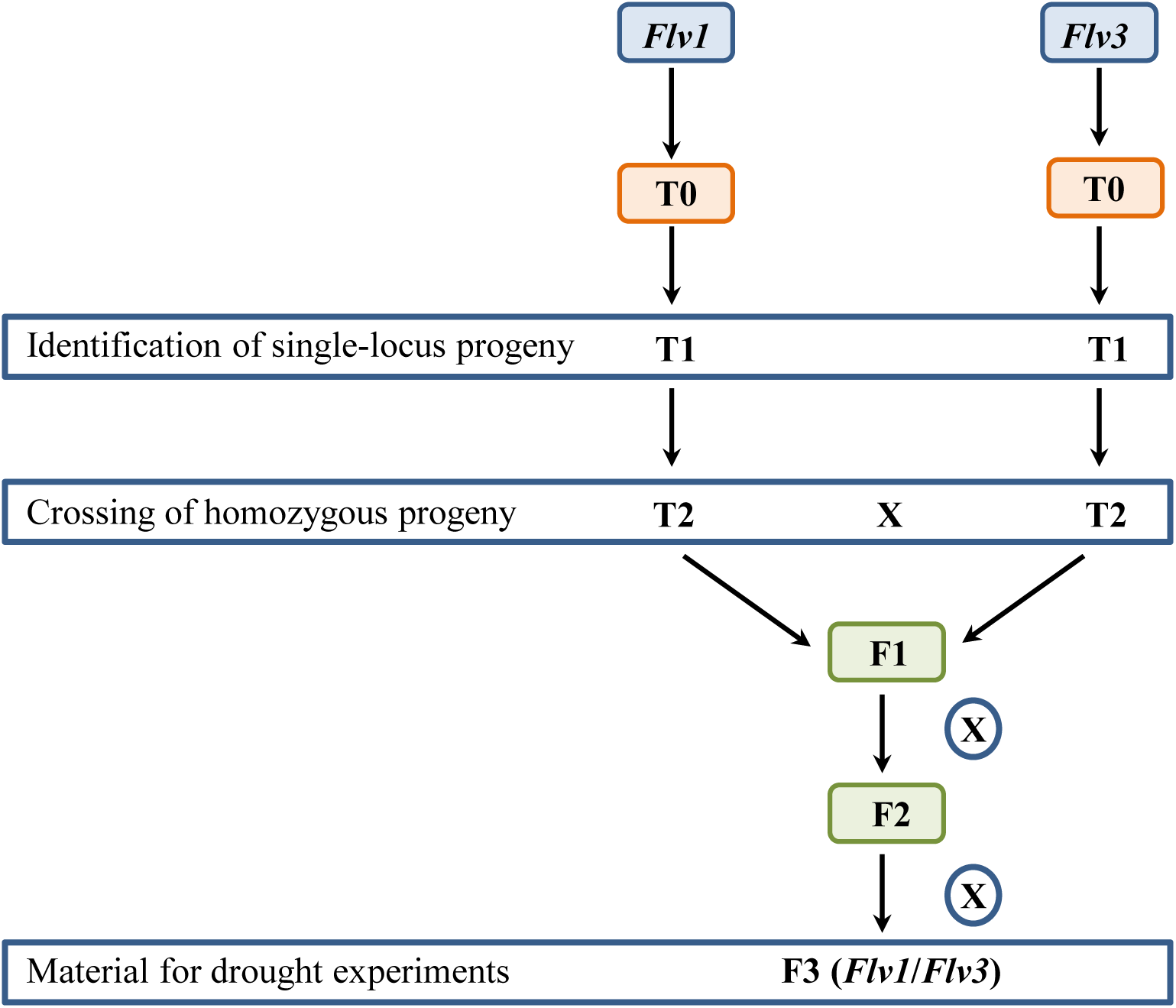
A model describing the steps for producing double-homozygous plants harbouring *Flv1/Flv3* to conduct drought experiments.

**FIGURE S3.**
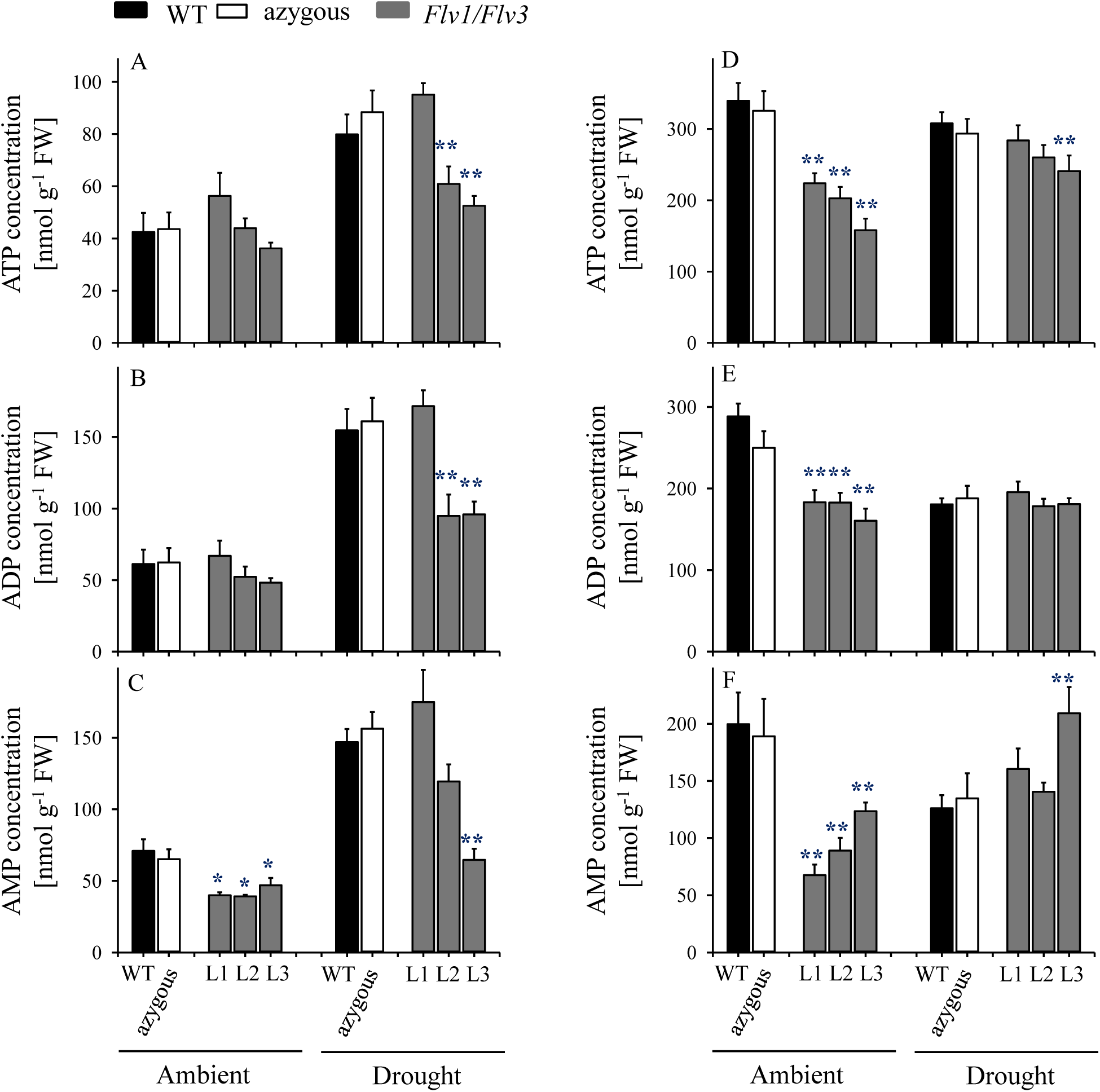
Influence of heterologously expressing *Flv* genes on adenosine nucleotides in the flag leaves of barley plants grown either under ambient conditions or exposed to drought stress at the seedling stage (A-C) and the reproductive stage (D-F). Adenine nucleotide levels were measured in the same samples used for carbohydrate determinations (Figure 5). (A, D) ATP, (B, E) ADP, (C, F) AMP. Lines L1-L3 harbour both *Flv1 and Flv3* genes. Data are shown as means ± SE (*n = 5-*6). **; *: means differed significantly (P ≤ 0.01 or P ≤ 0.05, respectively) from those of non-transgenic plants. FW, fresh weight.

**FIGURE S4.**
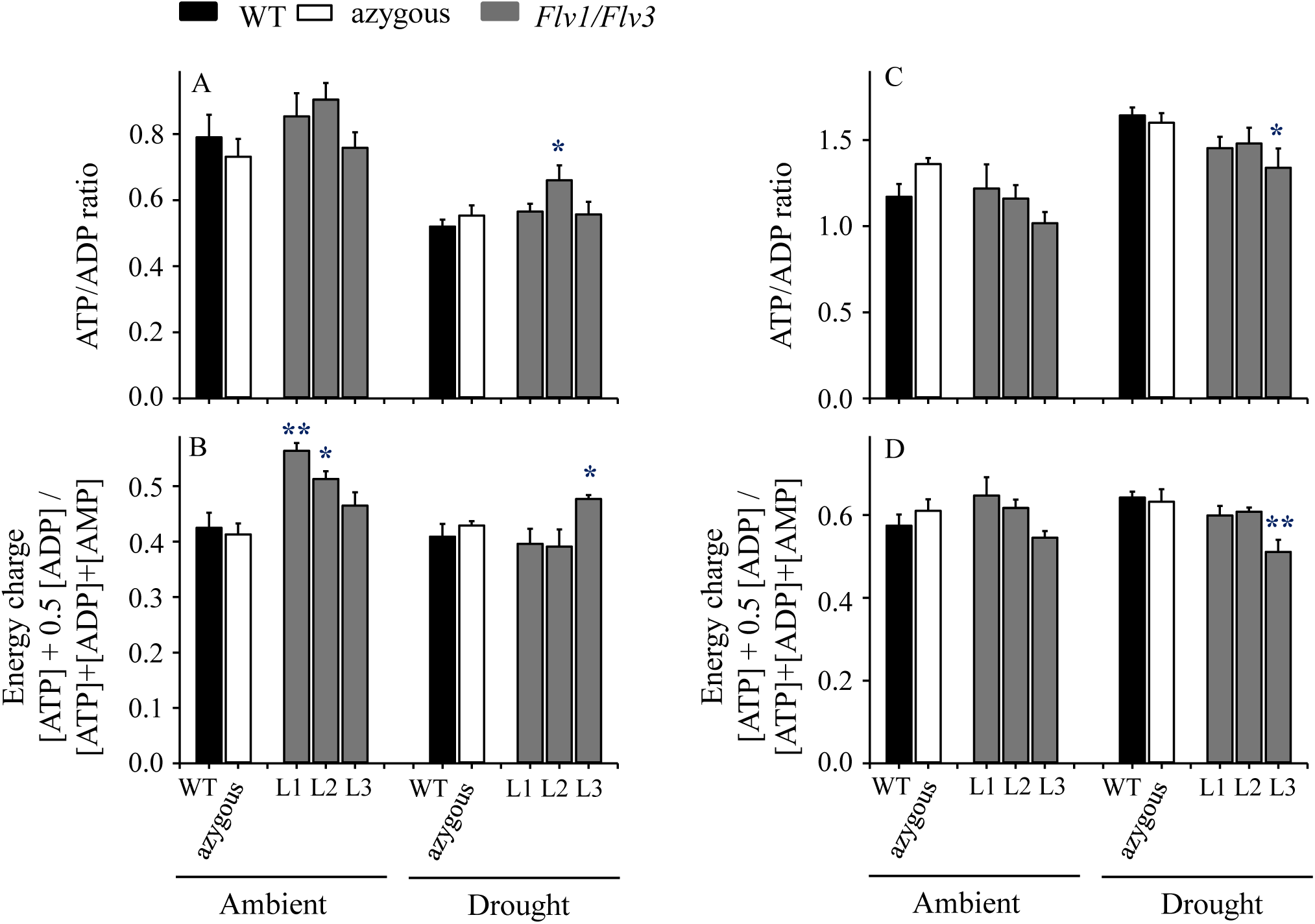
Effect of heterologously expressing *Flv* genes on the energy status of flag leaves of barley plants grown either under ambient conditions or exposed to drought stress at the seedling stage (A-B) and the reproductive stage (C-D). (A, C) ATP to ADP ratio, (B, D) energy charge. Lines L1-L3 co-express *Flv1 and Flv3* genes. Data are shown as means ± SE (*n = 5-*6). **; *: means differed significantly (P ≤ 0.01 or P ≤ 0.05, respectively) from the performance of non-transgenic plants.

## SUPPLEMENTARY TABLES

**Table S1** Sequence-specific forward (F) and reverse (R) primers, annealing temperature and extension time used for PCR and qRT-PCR analysis.

**Table S2** Effect of heterologously expressing *Flv* genes on amino acid contents in flag leaves of barley plants grown either under ambient conditions or exposed to drought stress at the vegetative stage. Lines L1-L3 harbour both *Flv1* and *Flv3* genes. Data are shown in nmol g^-1^ FW and as means ± SE (*n* = 6-7 for WT and azygous and *n* = 8-14 for transgenic lines). Yellow and blue shading: means differed significantly (P ≤ 0.01 and P ≤ 0.05, respectively) from those of non-transgenic plants. FW, fresh weight.

**Table S3** Effect of heterologously expressing *Flv* genes on amino acid contents in flag leaves of barley plants grown either under ambient conditions or exposed to drought stress at the reproductive stage. Lines L1-L3 harbour both *Flv1* and *Flv3* genes. Data are shown in nmol g^-1^ FW and as means ± SE (*n* = 6-7 for WT and azygous and n = 8-14 for transgenic lines). Yellow and blue shading: means differed significantly (P ≤ 0.01 and P ≤ 0.05, respectively) from those of non-transgenic plants. FW, fresh weight.

## Notes

### Competing Interest Statement

The authors have declared no competing interest.

